# Representational magnitude as a geometric signature of image and word memorability

**DOI:** 10.1101/2025.09.01.673067

**Authors:** David Amadeus Vogelsang, Micha Heilbron

## Abstract

What makes some stimuli more memorable than others? Recent work has shown that image memorability is predicted by the magnitude of population responses in both monkey inferotemporal cortex and convolutional neural networks, suggesting that stimuli that more strongly activate distributed feature representations are more likely to be remembered. However, it remains unclear whether this effect is specific to the visual domain or reflects a more general property of distributed representations. Here, we show that representational magnitude predicts memorability beyond the visual domain: the effect not only replicates for images in an independent dataset, but extends to lexical memory. Across three large-scale word datasets, word embedding magnitude reliably predicts recognition memorability, independent of word frequency, valence, or length. However, the effect did not extend to a recently published voice memorability dataset, possibly reflecting distinct features driving auditory memory. Together, the cross-domain generalization suggests that the representational magnitude effect is a general property of distributed representations. Memorability, on this view, is inherent to encoding: stimuli that activate more features, and activate them more strongly – whether in brains or artificial neural networks – leave a larger representational footprint, and therefore a more lasting memory trace.

## 1. Introduction

Why do we remember some stimuli better than others? Research on memory often takes a subject-specific approach, focusing on the cognitive processes behind memory and individual differences. This perspective is based on the idea that memory is highly personal and influenced by cognitive factors such as attention (Wakeland-Hart et al., 2022). However, while factors like experimental conditions and individual differences influence memory performance, research over the past decade has found that certain stimuli are inherently more memorable than others across participants (Bainbridge et al. (2013, 2017, 2019); Isola et al. (2013); Wakeland-Hart et al. (2022); Aka et al. (2023); Kramer et al. (2023); Revsine et al. (2025)). By analyzing aggregated task scores for each stimulus memorability can be quantified, with a high level of agreement be-tween participants across stimulus types and experimental contexts. This has been observed across images (Isola et al. (2013); Kramer et al. (2023)), words (Aka et al. (2023); Madan (2021)), voices (Revsine et al. (2025) and even in art work (Davis and Bainbridge (2023)) and dance moves (Ongchoco et al. (2023)). Further support to the idea that memorability is an intrinsic property of stimuli was provided by Needell and Bainbridge (2022), who showed that a dedicated deep neural network (DNN) model, specifically fine-tuned to predict human memorability, could accurately predict memorability (even on novel, out of sample stimuli) based solely on the features of an image.

Some studies have found that images that are more typical (e.g. how typical is this image of a banana?) are more memorable (Bainbridge and Rissman (2018); Bainbridge et al. (2019)), although other studies have found the opposite (Lukavský and Děchtěrenko (2017)) or found a more complex relationship between typicality and memorability (Kramer et al. (2023)). Remarkably, memorability not only varies consistently across humans but also in rhesus monkeys (Jaegle et al., 2019), suggesting memorability is independent of human-specific visual experience (e.g., vehicles, buildings etc.) and language ability.

Although the exact relationship between image content and memorability is still a matter of scientific scrutiny, the availability of image memorability scores has enabled research into the brain’s representational correlates of memorability and the development of DNN models that quantify and predict image memorability. Recently, a series of studies reported a dual coding scheme that predicts how well images will be discriminated and remembered (Jaegle et al. (2019); see Rust and Mehrpour (2020) and Rust and Jannuzi (2022) for reviews). Specifically, Jaegle and colleagues (2019) observed that whereas the angle of the response vector between two images quantifies how well they can be discriminated, the length of an image’s response vector (“L2 norm” or response magnitude), significantly predicted its inherent memorability, as quantified across a large image dataset (LaMem) and their memorability scores (Khosla et al. (2015)). Strikingly, this dual coding scheme was present in both activation patterns in rhesus macaque monkey inferotemporal cortex (IT), and in deep convolutional neural networks (CNNs) trained for object recognition (Jaegle et al. (2019)). Thus, it appears that brain-like representations of stimulus memorability emerge in artificial neural networks that are not explicitly trained to predict memorability. This coding scheme is theoretically significant because it suggests that memorability is not conferred onto a stimulus by downstream memory systems, but is inherent to how the stimulus is encoded in the first place. In this view, a stimulus that activates more features, and activates them more strongly, is more likely to be remembered – simply because of the magnitude of its initial encoding, not because subsequent memory systems treat it differently. Such an account aligns with cognitive (signal detection) memory strength models (Wixted, 2007), which formalise recognition memory performance as reflecting the strength of the underlying trace – here, it’s simply the magnitude of the initial representation that specifies what makes a memory trace strong. Consistent with this account, cognitive neuroscience research has shown that intrinsically memorable stimuli elicit stronger neural responses during encoding, resulting in more robust memory traces. In particular, larger neuronal response magnitudes during encoding have been linked to subsequent memory success (Vogelsang et al., 2018; Jaegle et al., 2019). If distributed representations in deep neural networks approximate high-dimensional feature spaces similar to those in the brain, then representational magnitude (operationalized as the L2 norm) may serve as a computational proxy for encoding strength or response magnitude, analogous to larger neural responses observed in biological systems. The fact that this emerges in both monkey visual cortex and convolutional neural networks underscores the alignment in how biological and artificial vision systems encode information.

However, these striking findings leave some questions unaddressed. First, it remains unclear whether the findings reported by Jaegle et al. (2019) can be replicated in other large datasets of image memorability. Second, and more importantly, it is unclear whether this coding scheme is inherent to visual perception and image memorability, or whether it reflects a more general principle, applicable to stimulus memorability (and stimulus encoding) more broadly. One possibility is that it is specific to visual memorability and visual representations because it derives from the representational convergence between the ventral visual stream and deep CNNs (DiCarlo and Cox (2007); Yamins and DiCarlo (2016)) – a neural network architecture that is, after all, explicitly modelled after the visual cortex (Lindsay (2021)). Alternatively, the emergent coding scheme (where L2 norm or representational magnitude predicts memorability) may be a more general principle of stimulus encoding in distributed representations, that would be applicable to all distributed representations, irrespective of stimulus modality or ‘brain-likeness’ of the network architecture used.

Here, we address both questions by first attempting to replicate the basic effects reported by Jaegle et al. (2019) to a different and independent large dataset of image memorability, spanning 26,107 naturalistic object images across 1,854 diverse categories tested in over 13,000 participants (THINGS; Hebart et al. (2020)). Secondly, we want to examine whether the same principle applies to different memorability domains, namely, word memorability. To test this, we analysed 3 datasets of word recognition memorability, covering a total of over 8500 memorability scores across 800 participants. To quantify representational magnitude, we computed the L2 norm of word embeddings, vector representations that captured a lot of information about words but are derived from neural networks that are not in any way inspired by or modelled after the human brain. Interestingly, we find that first, the L2 norm effect as reported by Jaegle et al. (2019) is robust and consistently replicates. Moreover, we find the exact same ‘representational magnitude effect’ for word recognition memorability, indicating that the effect is not specific to visual perception or sensory cortex, but reflects a more general property of stimulus encoding in distributed representations, perhaps related to the alignment between individual stimuli and dominant encoding ‘axes’ in both biological brains and neural networks. Finally, we evaluate the same effect in a recent voice memorability dataset covering a total of over 600 memorability scores across 2700 participants (Revsine et al. (2025)). However, this representational magnitude effect (using the L2 norm of waveform embeddings of Wav2vec) was not found in the voice memorability dataset, which could relate to particularities of this dataset or reflect limitations in the generality of the phenomenon.

## 2. Methods

### Datasets

We analysed the effect of representational magnitude on recognition memory in six large-scale datasets – one image, three word memorability datasets, and two voice memorability datasets – encompassing in total over 35,000 memorability ratings across visual, word and voice stimuli. We outline each in turn.

#### Kramer et al. (2023; THINGS)

Memorability scores were obtained from the THINGS dataset as reported in (Kramer et al. (2023)). Details on the study design can be found in Kramer et al. (2023) but we provide a brief summary here. A total of 13,946 participants from Amazon Mechanical Turk (AMT) completed a continuous recognition repetition detection task using 26,107 images from the THINGS database (Hebart et al. (2020); see https://things-initiative.org/). The THINGS database includes 1,854 object concepts spanning 27 categories, each with at least 12 exemplar images. Memorability scores were collected for all 26,107 images in the THINGS database by having participants view a stream of images and press a key when they recognized a repeated image. Each participant saw a unique pseudorandom subset of 187 images, with approximately 40 responses gathered per image. Memorability was calculated as the average hit rate (HR) across participants for each image, adjusted for the false alarm rate (FA), which accounts for the tendency to incorrectly recognize novel images as familiar (see figure 1 for illustration). This correction is typically applied per image to reflect variations in how images may evoke a sense of familiarity, even when they have not been seen before (Rust and Mehrpour (2020)). The raw data can be obtained here: https://osf.io/5a7z6/.

**Figure 1:**
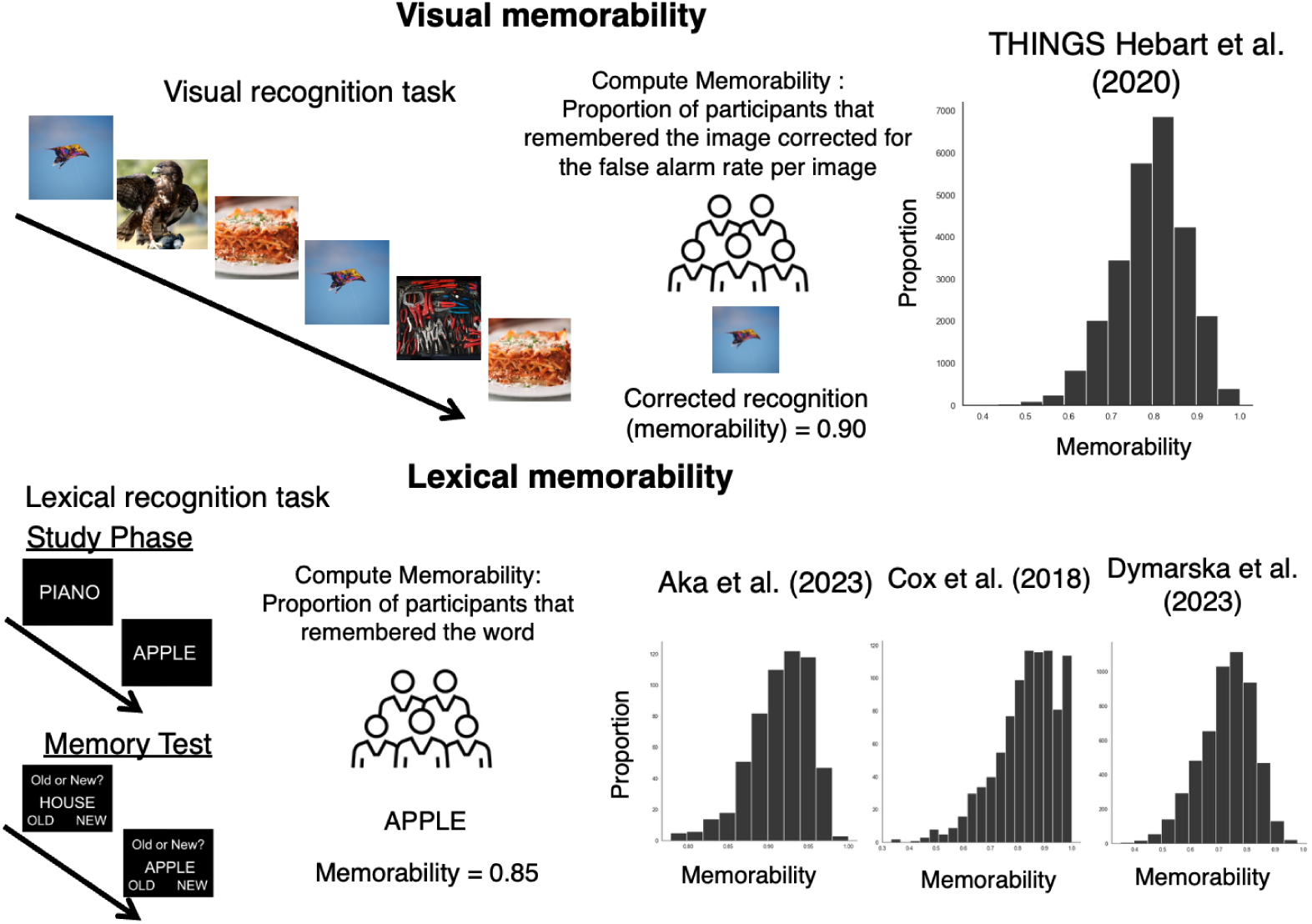
Overview of visual and word memorability paradigm. Images/words are studied and in a subsequent recognition memory test participants indicate whether a stimulus is old or new. For images, memorability is computed by calculating the proportion of participants that remembered a certain image corrected for the false alarm rate per image. For words, memorability is computed as the proportion of participants that correctly recognised a word in an old/new recognition memory test. Memorability score distributions are provided for the visual and word datasets in the graphs on the right.

#### Aka et al. (2023)

For the Aka dataset, we used the word recognition and recall memorability scores which were calculated on datasets from the PEERS database. A detailed description of the PEERS database has been described by Kahana et al. (2024) and further details of the memorability scores has been reported by Aka et al. (2023). In short, the Penn Electrophysiology of Encoding and Retrieval Study dataset (PEERS; available at http://memory.psych.upenn.edu/Data_Archive) consists of recognition and recall memory data. The recognition memory dataset includes 171 participants (ages 18–30) who contributed 3120 sessions. Each session involved studying 12–16 lists of 16 words (from a pool of 1638 words) and completing a recognition test with 320 probe words. The recall memory dataset includes 98 participants (ages 18–30) who contributed 2254 sessions. Each session involved studying 24 lists of 24 words (from a pool of 576 nouns), followed by a recall test after a brief arithmetic distraction task. Recognition and recall probabilities were estimated as the percentage of participants who correctly recognized/recalled each word (see figure 1 for illustration). The Aka et al. (2023) dataset consists of 576 words with recognition and recall memorability scores. The Aka et al. (2023) data is available here: https://osf.io/7phj6/files/osfstorage.

#### Cox et al. (2018)

We obtained recognition memorability scores from the Cox et al. (2018) dataset. Participants performed various memory tests (e.g. associative memory and cued recall test) of which only the recognition memory and the free recall were used in this study. During the study phase of each task, 453 participants viewed 20 word pairs, each presented for two seconds. In the single-item recognition memory task, they identified which of 20 words had appeared on the studied list, with 10 words serving as new ‘foils’. In the free recall task, participants attempted to recall as many words as possible from the study list. Cox et al. (2018) did not compute memorability scores themselves hence we computed recognition and recall memorability scores for each studied word by taking again the proportion of participants who correctly recognized/recalled each word. In total, 924 words with their corresponding recognition and recall memorability scores were computed in this dataset. The raw Cox et al. (2018) data files can be obtained here: https://osf.io/dd8kp/.

#### Madan (2021)

Madan (2021) only reported the recall memorability scores that were obtained from the Penn Electrophysiology of Encoding and Retrieval Study (PEERS), and therefore we are only using this dataset for the free recall memorability analysis as recognition memorability data was not available in the Madan dataset. In each session, participants studied 12–16 lists of 16 words, with each word displayed for 3 seconds followed by a short interval. After the final word, a brief delay preceded a free recall test, where participants had 75 seconds to vocally recall as many words as possible. Recall data were examined from 147 young adults (ages 16–30), each completing 20 sessions in the PEERS experiments. The word pool consisted of 1,638 words from the University of South Florida free association norms database, selected for their semantic relatedness and suitability for size and animacy judgments. In total, 1638 words and their corresponding recall memorability scores were used in the Madan dataset. The raw datafiles can be obtained here: https://osf.io/spqjz/

#### Dymarska et al. (2023)

We used the data as reported by Dymarska et al. (2023) who analysed data from two large recognition memory studies for words (over 5300 words) as reported by Cortese and colleagues (Cortese et al. (2010, 2015)). We provide a brief summary of the Cortese studies here: Across two large studies, 207 participants (87 in Cortese et al. (2010); and 120 in Cortese et al. (2015)) were presented with 5301 words that were used a recognition memory test. In both studies, participants studied lists of 50 words at a time in preparation for a later recognition task. This task required them to make an old/new judgment on each presented word, with half of the target words being old and the other half new. Hit rate, false alarm rate and corrected hit rates per stimulus (i.e. word) were computed, of which for the purpose of this current study we only used the hit rate (i.e. memorability score) per word to make it comparable to the other word datasets used in the current study. The raw data used in the Dymarska study can be downloaded here: https://osf.io/r8fmb/.

#### Revsine et al (2025)

We used the voice memorability data collected by Revsine and colleagues (2025), which comprised of three voice memorability experiments. In their study, 3,340 Amazon Mechanical Turk participants completed a 10-minute continuous recognition memory task, where they listened to the same sentence (“She had your dark suit in greasy wash water all year” in experiment 1, and “Don’t ask me to carry an oily rag like that” in experiment 2) being spoken by different speakers and pressed a key when they heard a repeated speaker. Stimuli were drawn from the TIMIT speech corpus. Memorability (measured with d’) was collected for each voice clip in Experiments 1 and 2. All raw data can be downloaded here: https://osf.io/pybwd/.

### Data Analysis

An overview of our analysis approach is provided in figure 2. We computed the correlation (i.e. Spearman correlation) between the response magnitude (after the ReLu activation function) at all layers of a convolutional neural network (i.e. AlexNet) and memorability scores for images in the THINGS dataset. We then investigated whether we could extend this analysis to the domain of word memorability by examining the correlation between the response magnitude of a simple neural network (i.e. Word2vec) and recognition and recall memorability scores for words. For the images in the THINGS dataset, we used the memorability scores as reported by Kramer et al. (2023). For each image in the THINGS dataset, we extracted the features of each layer in AlexNet. AlexNet is a deep convolutional neural network (CNN) designed for image recognition, and it contains eight layers of which five are convolutional and three are fully connected (Krizhevsky et al. (2017)). The response magnitude (L2 norm) for each layer was calculated using the entire output vector of the hidden layer. For every layer, we calculated Spearman correlation coefficients for all rectified linear unit (ReLu) layers. Correlations between the L2norm of each layer and memorability scores were computed across the entire set of 26,107 images from the THINGS dataset.

**Figure 2:**
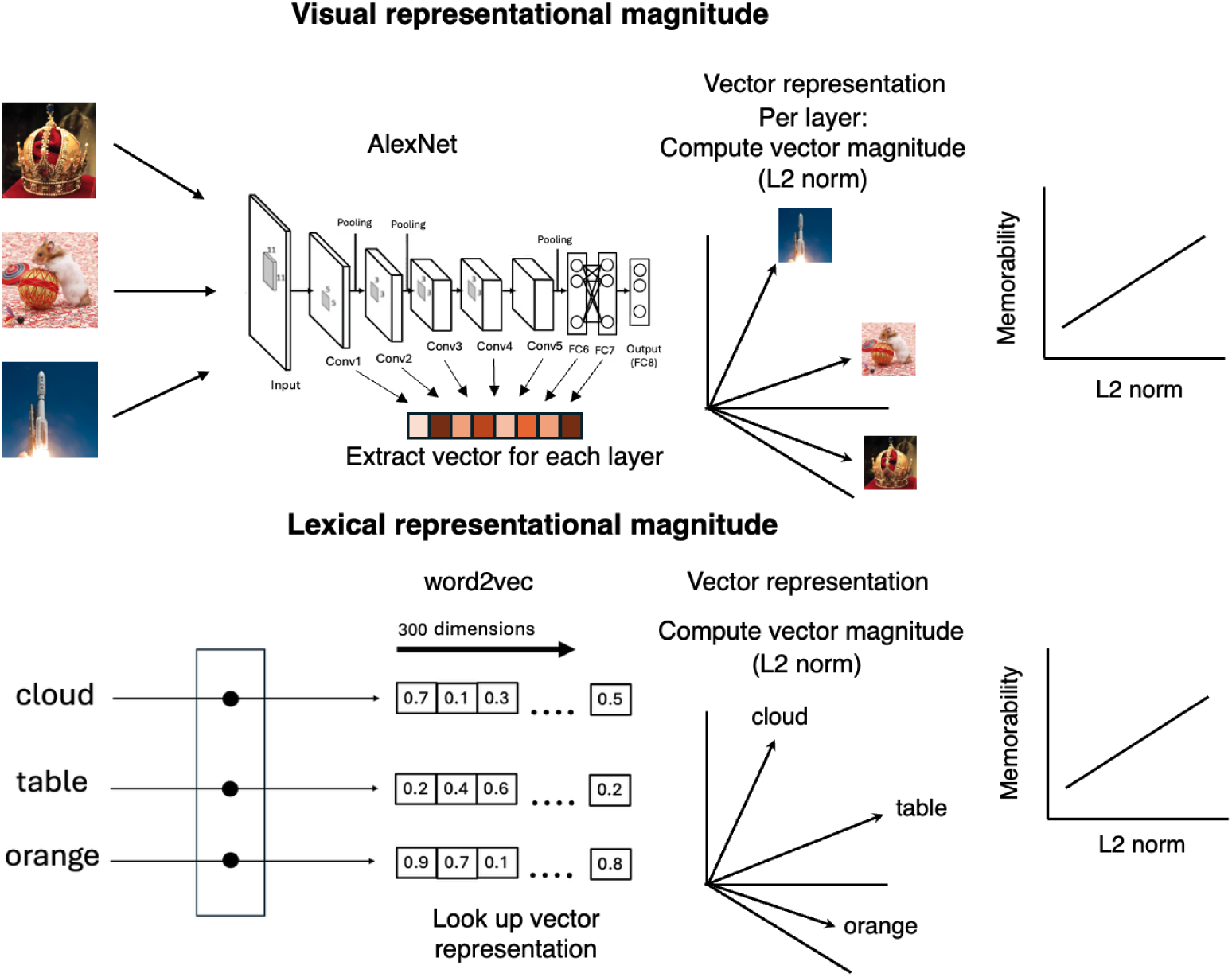
For each image in the THINGS dataset, we extracted the features (i.e. vector) representation of each layer of AlexNet and computed the vector magnitude (i.e. L2 norm) of each layer. Finally, we correlated the per image memorability score with the vector magnitude. For words, we extracted for each word the vector representation of the 300-word embeddings from Word2vec. We again computed the L2 norm of each vector representation and correlated this with the recognition memorability score per word.

For the word memorability analysis, we utilized the pretrained Word2vec model ’GoogleNews-vectors-negative300’ (Mikolov et al. (2013)). Word2vec is a neural network-based model that learns vector representations of words by analysing their usage in large text corpora. It captures semantic relationships by positioning words with similar meanings closer in a high-dimensional space (Mikolov et al., 2013). The model uses the Skip-gram architecture, which efficiently learns high-quality word vectors that capture a wide range of precise syntactic and semantic relationships and is therefore good at predicting surrounding words given a target word (Mikolov et al. (2013)). This model was trained by Google on approximately 100 billion words from the Google News dataset using the skip-gram architecture (Mikolov et al. (2013)). Here, we extracted for each word in each dataset (i.e. Aka, Cox and Madan) the 300-word embeddings and computed the L2 norm. As a final step, we computed the Spearman correlation between the L2 norm of each word and their corresponding recall/recognition memorability scores.

For the voice memorability analysis, we used Wav2vec, which is a self-supervised deep learning model developed for speech representation learning from raw audio waveforms (Baevski et al. (2020)). Unlike traditional approaches that rely on handcrafted acoustic features, Wav2vec learns feature representations directly from unlabeled audio data by predicting masked segments of the input waveform based on surrounding context. Its architecture consists of a multi-layer convolutional feature encoder that processes the raw waveform, Transformer-based context network that captures long-range temporal dependencies. This approach enables the model to capture phonetic and prosodic information that can be fine-tuned for various downstream tasks, including speech recognition. Importantly, although Wav2vec was developed for speech recognition, its representations are constructed from the raw audio and should capture speaker-specific information very well (Fan et al., 2021). Its general purpose representations make the network well-suited to test the representational magnitude effect in the Revsine dataset. For each audiofile in the Revsine dataset (who used the TIMIT database), we extracted the features from all Wav2vec layers and computed the L2 norm. We computed the d’prime score, which Revsine used as their measure of memorability, for each audiofile and conducted a Spearman correlation between the L2 norm and memorability scores.

To estimate the reliability of the observed correlation between representational magnitude and memorability scores, we employed a non-parametric bootstrapping procedure. Specifically, we computed the Spearman correlation coefficient between the two variables and then generated a 95% confidence interval via resampling. This was done by repeatedly sampling the dataset with replacement 10,000 times, preserving the pairing between each data point (i.e., paired resampling of the two variables). For each resampled dataset, we recalculated the correlation coefficient, resulting in a distribution of 10,000 correlation values. The 2.5th and 97.5th percentiles of this distribution were taken as the lower and upper bounds of the confidence interval. This approach allowed us to quantify the uncertainty in the estimated correlation without relying on assumptions of normality or homoscedasticity.

### Control Analysis

In our main analyses, we examined the relationship between vector magnitude (L2 norm) and memorability. However, one potential alternative interpretation is that the observed relationship between L2 norm and memorability may be driven by confounding factors, such as image typicality or word frequency, valence or size. If higher L2norm values merely reflect more typical images or more frequently used words, it could be that the effect attributed to representational magnitude could instead be explained by image typicality/word frequency. To address this possibility, we conducted a a set of control analysis. Specifically, for images in the THINGS dataset, Kramer et al. (2023) computed typicality based on both the 49 objects dimensions as obtained and described in Hebart et al. (2020) as well as from a deep convolutional neural network (DNN). In the 49 dimension object feature space, items that are closely clustered within a concept (i.e. within the concept of airplane, an image of an airplane is more similar to other images of airplanes) are the most prototypical, whereas those that are more widely spaced are the most atypical (e.g. an image of an airplane that looks very different from the other images of airplanes). To compute typicality from the DNN, Kramer et al. (2023) calculated the average similarity (Pearson correlation) between each image and all other images representing the same concept (e.g., the similarity of a given strawberry image to all other strawberry images in the THINGS dataset). This procedure was repeated across all layers of the DNN, yielding a typicality score for each layer. For the purpose of our regression analysis, we take those typicality values of the last layer of the DNN. The typicality values of the object space and DNN typicality were provided by the authors and can be obtained through the following link: https://osf.io/5a7z6/files/osfstorage. To examine possible confounding variables on the L2 norm effect on word memorability, we computed a Spearman correlation matrix of the most prominent word features that were provided in the Aka et al. (2023) dataset (see figure 7A). To identify candidate confounding variables, we examined the relationships between L2 norm and available lexical features and selected those showing substantial associations for inclusion in the control analyses. As can be seen in the correlation matrix, three word features showed a significant correlation in the same direction as the L2 norm of Word2vec, namely, word frequency, valence and size. These three features could therefore be a confounding variable explaining the positive association between the L2 norm and recognition memorability. To investigate the effect of frequency on the L2 norm effect, we obtained frequency ratings per word using the SUBTLEX, which calculated word frequency based on television and film subtitles (Brysbaert and New (2009)). Valence and size ratings were obtained from the Aka et al. (2023) dataset.

To ensure that the L2norm effect on image and word memorability was not confounded by image typicality or word frequency, valence or size, we conducted partial correlation analyses. Specifically, we assessed the relationship between representational magnitude (L2 norm) and memorability while statistically controlling for potential confounding variables. This was implemented by regressing out the confounding variables from both L2 norm and memorability using ordinary least squares (OLS) regression, and subsequently correlating the resulting residuals. This approach yields partial correlation coefficients that reflect the association between L2 norm and memorability independent of typicality (for images) and frequency, valence, and size (for words), allowing for direct comparison with the original zero-order correlations.

## 3. Results

In the primary analysis, we assess the effect of representational magnitude on recognition memory – as reported by Jaegle et al. (2019) – in 4 independent datasets, one of image memorability, three of word recognition memorability (see Figure 1). We first asked whether we could replicate the effect reported by Jaegle et al. (2019) of visual representational magnitude (based on CNN embeddings) on visual recognition memory in a different dataset. For this we turn to the THINGS-memorability dataset (Kramer et al. (2023)), which is a dataset of 26,107 images (see Methods). We computed the L2norm of each layer in AlexNet and correlated (Spearman correlation) it with the memorability scores of each of the 26107 images in the THINGS dataset. We ran this analysis for all 7 layers of the network. The results are displayed in Figure 3. As can be seen, the correlation between the L2norm and memorability is not significantly different from zero in early layers of the network but then increases significantly in later layers of the network (Layer 1: r = -.018, p = 0.004, 95% CI [-.03, -.006]; layer 2: r = .003, p = .62, 95% CI [-.009, .015]; layer 3: r = .007, p = .29, 95% CI [-.006, .019]; layer 4: r = .041, p = 2.5 e-11, 95% CI [.029, .053]; layer 5: r = .038, p = 7.9e-10, 95% CI [.026, .050]; layer 6: r = 0.05, p = 2.6e-16, 95% CI [.038, .061]; layer 7: r = .057, p = 2.7e-20, 95% CI [.045, 069] (see figure 3B)). This pattern of no/low correlation between the L2norm and image memorability in the early layers of a convolutional neural network and a higher and significant correlation in later layers of the network is a replication of the findings reported by Jaegle et al. (2019). However, the highest correlations we observe in the later CNN layers (r=0.057) are lower in magnitude than those reported by Jaegle (r=.30), which may be attributed to our dataset being derived from a large online crowd-sourced experiment that is inherently more noisy.

**Figure 3:**
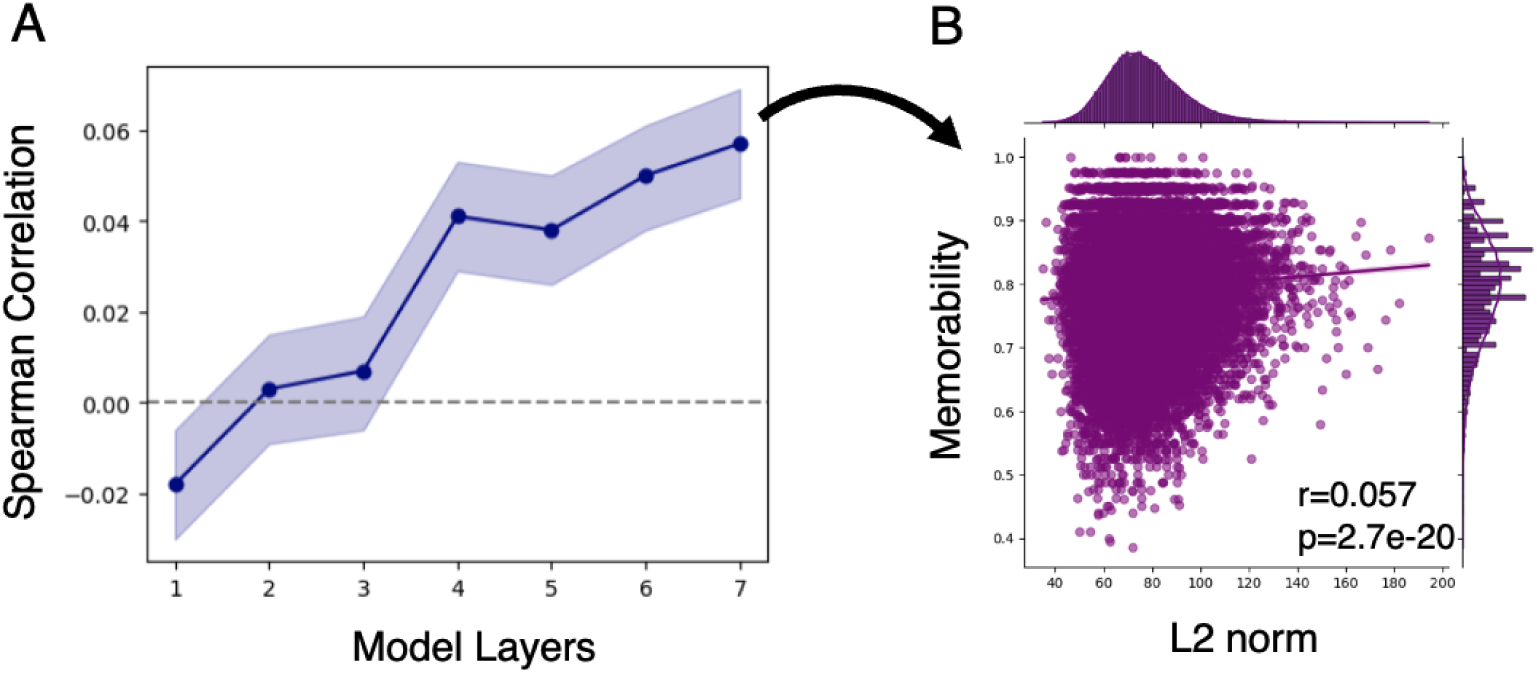
A) Spearman correlations for each layer of AlexNet (5 convolutional, 2 fully connected) between the magnitude of the representation at that layer (L2 norm), and the memo-rability score of each image in the THINGS dataset. B) Spearman correlation plotted for the last ReLU layer.

**Figure 4:**
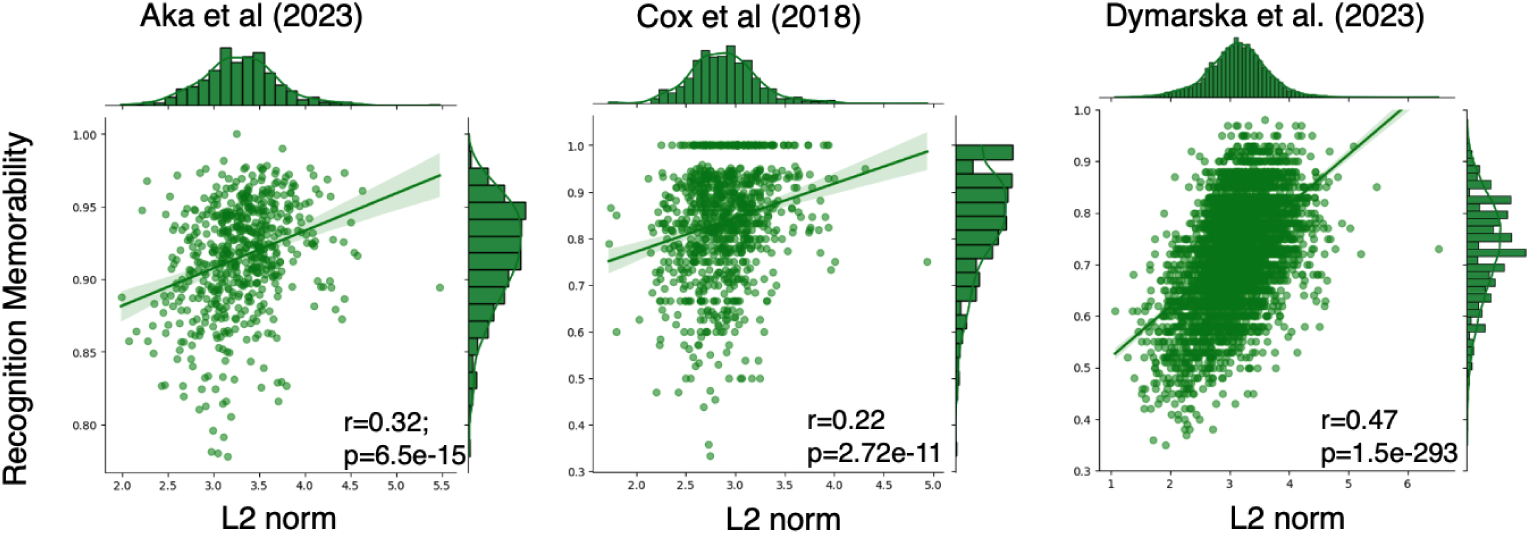
Spearman correlations between the L2norm and recognition memorability of each word across the three datasets.

After confirming that the effect reported by Jaegle and colleagues (2019) indeed replicates, we next asked whether this effect is specific to visual recognition memory or whether it applies to other domains, such as word and voice memorability, as well. For this, we first turned to the non-sensory domain of verbal recognition memory, for which we analysed three datasets, encompassing 8500 memorability scores across over 800 participants in total. In these datasets, participants performed a very similar recognition memory task, but memorised words instead of objects (see Figure 1 and Methods). We computed the L2 norm of the static word embeddings from Word2vec (see methods for details) for all words in every dataset and computed a Spearman correlation with the corresponding recognition memorability scores. As visible in Figure 3, this revealed a highly consistent and strong correlation for the Aka et al. (r = .32, p = 6.5e-15, 95% CI [.24, .39]), the Cox et al. (r = .22, p = 2.72e-11, 95% CI [.16, .28]) and Dymarska et al. datasets (r=0.47, p=1.5e-293, 95% CI [.45, .49]) between the embeddings representational magnitude and its recognition memorability. This result indicates that the L2norm effect as observed by Jaegle et al. (2019) may reflect a more general principle of stimulus encoding in distributed representations rather than a specific attribute of visual CNNs or visual cortex.

### Control Analysis

Given that embeddings are not inherently interpretable but encode a lot of stimulus information implicitly, it could be that the vector magnitude simply correlates with another stimulus attribute that is already known to co-vary with memorability. To rule this out, we consider several likely alternative explanations. For images, we considered the role of typicality. First, more “typical” object instances (e.g. in the visual case, typical cats) might be more similar to the instances seen during training (ImageNet), and hence more aligned to the feature detectors in AlexNet, leading to higher vector magnitudes. At the same time, typical images have also been found to be more memorable, although this relationship is complex and may depend on category-level expectations, distinctiveness and familiarity (Kramer et al. (2023)). To address this potential confound, we computed partial correlations between the vector magnitude of the last ReLu layer of AlexNet and image memorability while controlling for two different model-based image-typicality estimates (see methods for details). This was done by correlating the residuals of both variables after regressing out typicality. The results are displayed in figure 5B. For both object typicality (r = 0.062, p = 2.202e-23, 95% CI [0.05, 0.074]) and DNN typicality (r = 0.060, p = 3.825e-22, 95% CI [0.048, 0.072]), the relationship between L2 norm and memorability remained positive and statistically significant. This indicates that the effect of representational magnitude cannot be reduced to image typicality.

**Figure 5:**
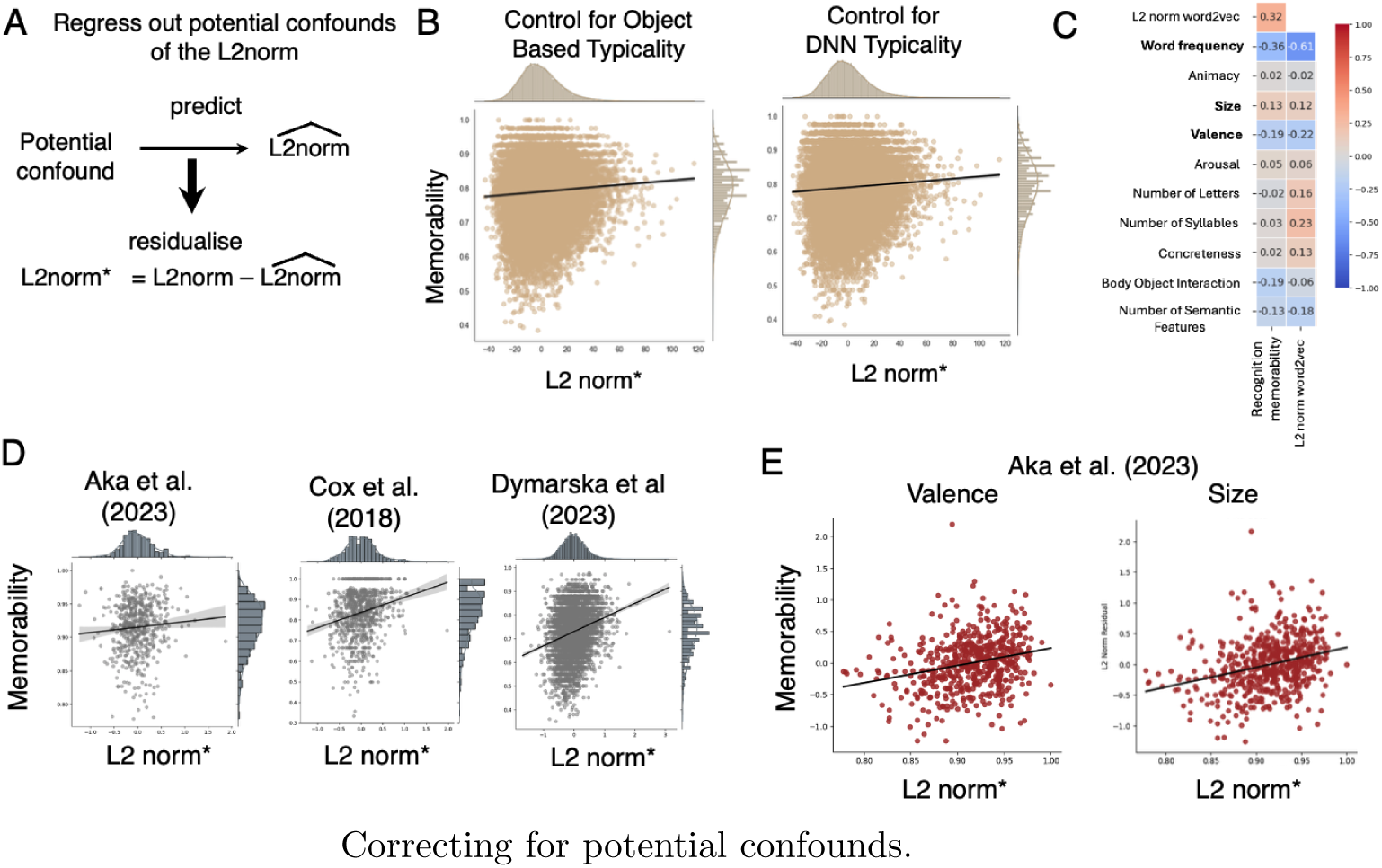
A) Regressing out image typicality (both object and CNN based) of the L2 norm B) Regressing out object and CNN based typicality, we still find a positive relationship between the L2 norm and image memorability. C) Correlation matrix of word features from the Aka dataset reveals that word frequency, valence and size show a similar significant correlation patterns as the Word2vec L2 norm. D)Regressing out word frequency, we will find a positive relationship between the L2 norm and recognition memorability in the Cox et al. (2018) and Dymarska et al. (2023) datasets, and a marginally positive effect in the Aka et al. (2023) dataset. E) Regressing out valence and size in the Aka dataset (only dataset with these features available) still revealed a positive relationship between L2 norm and recognition memorability.

For words, we first identified potential confounding features by examining which word features reveal a similar correlation pattern with recognition memorability as the L2 norm (see figure 5C). In short, a potential confounding feature has to either correlate positively L2 norm and/or memorability, or correlate negatively with both. As can be seen, word frequency, valence and size show correlations in the same direction as the L2 norm thereby revealing the possible confound that the L2 norm effect reflect a word frequency, valence or size effect in disguise. To address this potential confound, we conducted a control analysis in which we examined whether the L2 norm effect is still observed when accounting for word frequency, valence and size in lexical stimuli.

For word recognition memorability,we computed partial correlations between vector magnitude (L2 norm) and recognition memorability while controlling for word frequency. This was done by correlating the residuals of both variables after regressing out frequency. Across all three datasets, the relationship be-tween L2 norm and memorability remained positive after controlling for word frequency: Cox (r = 0.217, p = 2.665e-11, 95% CI [0.155, 0.278]) and Dymarska (r = 0.271, p = 6.151e-91, 95% CI [0.246, 0.296]) showed significant positive effects, whereas for the Aka dataset the effect was attenuated and reached trend-level significance (r = 0.079, p = .059, 95% CI [-0.003, 0.159], see Figure 5D). Second, word-level variables such as valence and size were only available for all items in the Aka dataset, as the Cox and Dymarska datasets included a broader range of word types beyond nouns. As a result, we were only able to control for valence and size in the Aka dataset. Importantly, even after statistically controlling for valence (r = 0.256, p = 4.496e-10, 95% CI [0.178, 0.331]) and size (r = 0.286, p = 2.668e-12, 95% CI [0.209, 0.360]), the relationship between L2 norm and recognition memorability remained strong and statistically significant (see figure 5E). All together, these results indicate that for word stimuli, the relationship between L2 norm and memorability are unlikely to be explained by word frequency, valence or size.

### Word Recall Memorability

In verbal memory experiments, it is common to assess memory either through recognition memory or free recall. Jaegle et al. (2019) were specifically interested in recognition memory: they found their original effect in a continuous recognition task, and linked their effect to neural representations in the ventral visual cortex during object recognition. However, since we found the effect is not specific to visual object recognition or visual processing, a natural question is to ask if it is specific to recognition memory in the first place, or whether it also extends to memory (i.e. “free”) recall. To address this, we repeated the same analysis, now analyzing the recall memorability scores in all three verbal datasets. The results are displayed in Figure 6. We found no significant correlation between the L2norm and recall memorability for the Aka (r = -0.017; p = .69, 95% CI [-.10, 0.07]) and Madan (r = .002, p = .92, 95% CI [-.046, .051]) datasets, however in the Cox dataset there was a significant positive correlation (r = .22, p = 7.1e-12, 95% CI [.16, .28]). These results reveal that the L2 norm effect as observed for both visual and word memorability does not extend convincingly to recall memorability scores. We discuss the implications of these findings in the discussion.

**Figure 6:**
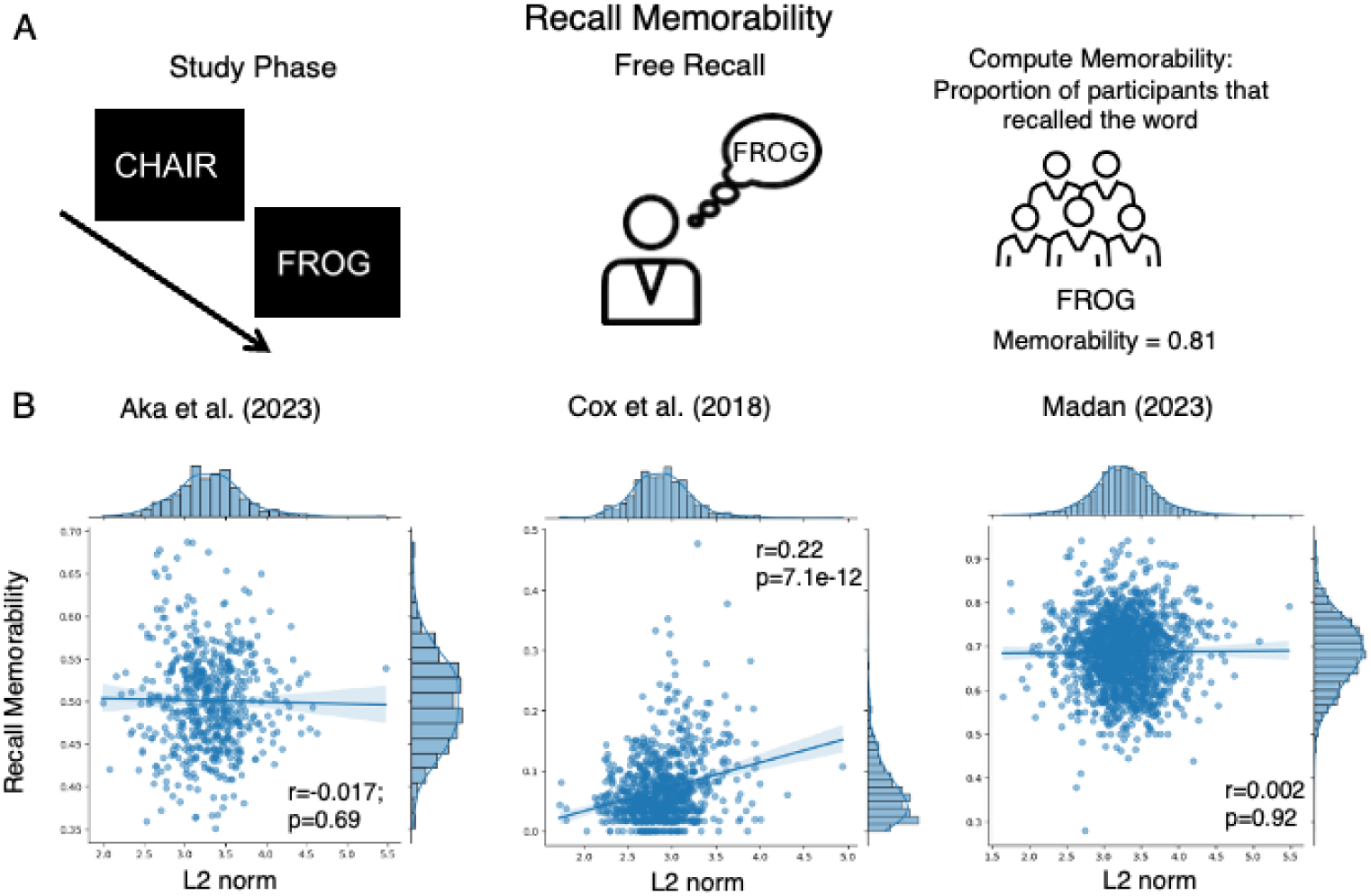
A) Recall memorability for words can be computed by calculating the proportion of participants that recalled a word. B) For the Aka et al. (2023) and Madan (2023) no significant correlation between the L2norm and memorability was found, however for the Cox et al. (2018) dataset there was a significant correlation.

**Figure 7:**
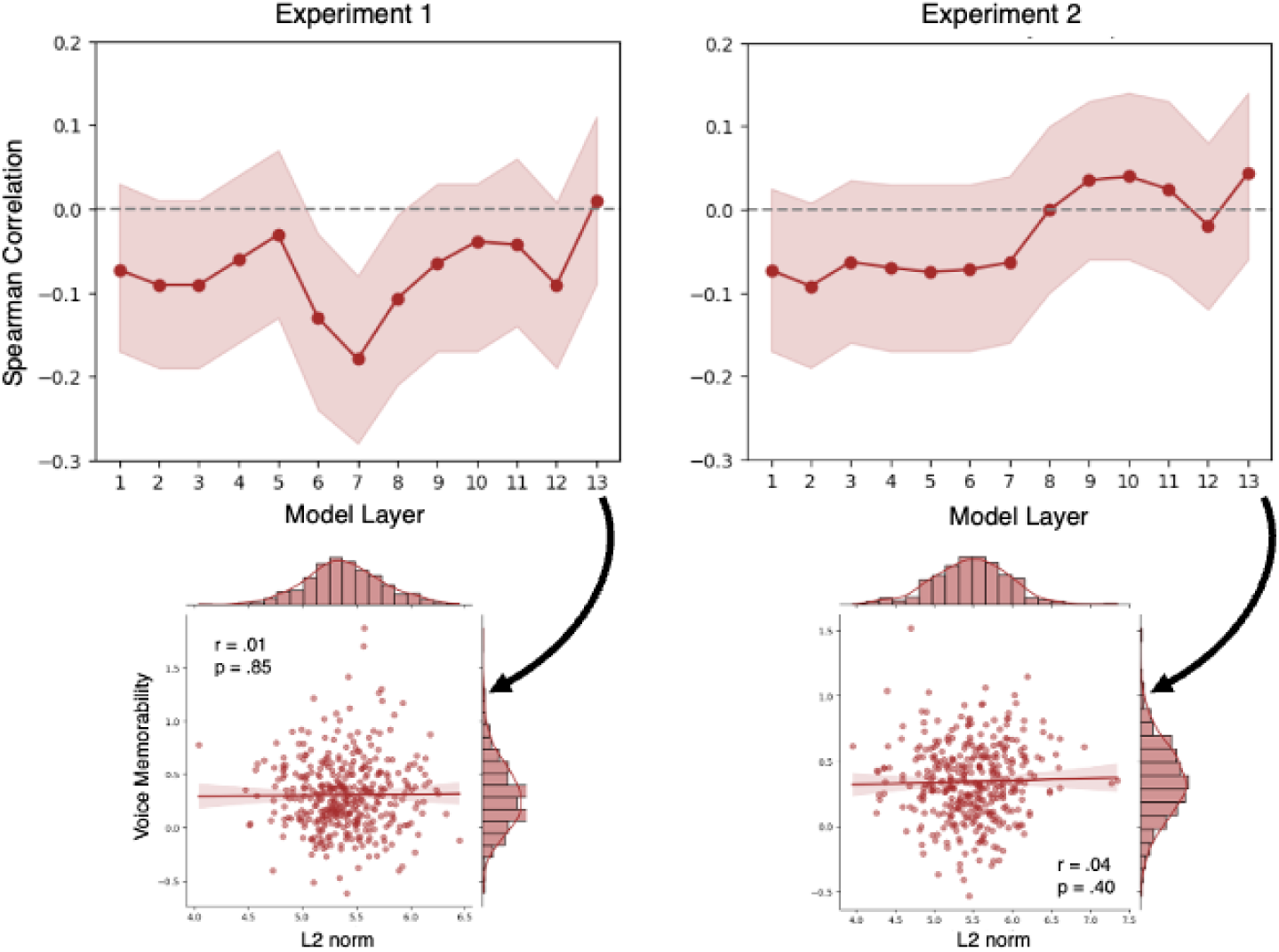
For both experiment 1 and 2 of the Revsine voice memorability datasets there was no reliable relationship between the L2 norm of any layer of wav2vec and voice memorability.

### Voice Memorability

After confirming that the vector magnitude and memorability relationship in the visual domain extends to the domain of word recognition memorability, we investigated whether this L2 norm can also be extended to the auditory domain of voice memorability. We analysed data from the first two experiments as reported by a recently published paper by Revsine et al. (2025), covering more than 600 voice memorability scores across over 2700 participants. We computed the L2 norm of all layers of Wav2vec (Baevski et al. (2020)) and ran across the voice stimuli a Spearman correlation between the L2 norm and the corresponding memorability score (see methods for details). The results are displayed in figure 7 and all statistical results are reported in table 1. In contrast to the visual and word memorability domain, for Wav2vec we did not observe a consistent significant relationship between the vector magnitude and memorability across experiment 1 and 2.

**Table 1:** Correlation between wav2vec vector magnitude and memorability across layers in experiment 1 and 2 of Revsine et al. (2025)

| Layer | Experiment 1 |  |  | Experiment 2 |  |  |
| --- | --- | --- | --- | --- | --- | --- |
|  | r | p-value | 95% CI | r | p-value | 95% CI |
| 1 | -0.07 | .15 | [-.17, .03] | -0.07 | .16 | [-.17, .025] |
| 2 | -0.09 | .08 | [-.19, .01] | -0.09 | .07 | [-.19, .008] |
| 3 | -0.09 | .08 | [-.19, .01] | -0.06 | .22 | [-.16, .035] |
| 4 | -0.06 | .23 | [-.16, .04] | -0.07 | .18 | [-.17, .03] |
| 5 | -0.03 | .54 | [-.13, .07] | -0.08 | .14 | [-.17, .03] |
| 6 | -0.13 | .009 | [-.24, -.03] | -0.07 | .16 | [-.17, .03] |
| 7 | -0.18 | .0004 | [-.28, -.08] | -0.06 | .22 | [-.16, .04] |
| 8 | -0.11 | .04 | [-.21, -.007] | -0.0008 | .99 | [-.1, .1] |
| 9 | -0.07 | .21 | [-.17, .03] | 0.04 | .49 | [-.06, .13] |
| 10 | -0.04 | .44 | [-.17, .03] | 0.04 | .44 | [-.06, .14] |
| 11 | -0.043 | .40 | [-.14, .06] | 0.02 | .64 | [-.08, .13] |
| 12 | -0.09 | .07 | [-.19, .008] | -0.02 | .71 | [-.12, .08] |
| 13 | 0.01 | .85 | [-.09, .11] | 0.04 | .40 | [-.06, .14] |

## 4. Discussion

In this paper, we investigated the relationship between the magnitude of DNN representations and memorability scores of linguistic and visual stimuli. We first replicated the findings of Jaegle et al. (2019), demonstrating that vector magnitude in later layers of a CNN (AlexNet) positively correlates with image memorability scores from the THINGS dataset, even after controlling for potential confounds such as image typicality. Extending this line of research to the linguistic domain, we found that the L2 norm of neural word embeddings was also positively correlated with recognition memorability across three independent datasets, demonstrating that the effect generalizes beyond visual recognition to the lexical domain. When we tested this effect in the auditory domain using voice memorability data from Revsine et al. (2025) and embeddings from Wav2vec, however, we did not observe a significant relationship between vector magnitude and voice memorability. Together, these findings provide a new perspective on DNN stimulus encoding and stimulus memorability across sensory modalities. The presence of the L2 norm effect in both visual and lexical domains suggests that it may reflect a domain-general property of representational strength, rather than a mechanism restricted to vision. More broadly, this highlights the potential of representational geometry as a computational framework for studying how intrinsic stimulus properties shape memorability across domains.

What does the representational magnitude effect reflect? We propose that representational magnitude captures how extensively a stimulus activates the features through which it is encoded. A stimulus that projects strongly onto many learned feature dimensions – in biological brain or a DNN representation – will have a larger L2 norm. In other words, it leaves a larger footprint in representational space, and thereby a stronger memory trace. In biological brains, stimuli that are subsequently remembered better tend to elicit larger neuronal amplitudes during encoding (Wagner et al., 1998; Brewer et al., 1998; Vogelsang et al., 2018; Jaegle et al., 2019), consistent with more extensive feature activation and the formation of stronger memory traces. From the perspective of cognitive theories of memory, this aligns with (signal detection) strength-based accounts (Wixted, 2007), which propose that variability in memory performance reflects differences in strength of the memory signal. Artificial neural networks provide a computational instantiation of this principle. In these systems, stimuli that activate learned features more strongly will have a larger magnitude DNN representation but will also – assuming sufficient alignment between the features encoded in DNNs and those encoded in biological systems (Chen and Bonner, 2025) – evoke stronger feature activation in biological brains, and therefore leave a stronger memory trace. On this account, the representational magnitude at encoding does not merely correlate with memory; it simply constitutes the strength of the memory trace. Importantly, this is not about activating more features indiscriminately, but about activating *relevant* features: for images, replicating Jaegle et al. (2019), we find that the effect is weak in early CNN layers but pronounced in later layers encoding higher-level semantic features – paralleling neuroimaging findings that memorability effects emerge in higher-order regions rather than early visual areas (Bainbridge et al., 2017).

Critically, this interpretation connects our neural network-based approach to classical computational models of recognition memory. In summed-similarity frameworks such as SAM, MINERVA, and GCM (Gillund and Shiffrin, 1984; Hintzman, 1988; Nosofsky, 1991), the recognition signal is computed by summing the similarity between a test probe and all stored memory traces—including the probe’s similarity to its own trace, or ‘self-similarity’. Traditionally, these models used distance-based similarity metrics in which self-similarity is constant across items, requiring additional mechanisms (such as distinctiveness ratings) to explain variation in hit rates (Nosofsky et al., 2025). However, when similarity is computed via dot product – as is natural for neural network representations – self-similarity becomes proportional to squared vector magnitude. From this perspective, the representational magnitude effect we observe in neural networks may provide a mechanistic basis for what classical memory models captured through auxiliary assumptions: items with larger representational magnitude automatically generate a stronger self-match during recognition, and therefore a stronger memory signal.

Our interpretation can also account for other, seemingly unrelated observations. For instance, Tuckute et al. (2025) found that words with a strictly one-to-one relationship with their meanings – having few synonyms and less semantic ambiguity (polysemy) – are more memorable. This aligns with such words having larger representational magnitudes. Indeed, the inverse relationship between polysemy and representational magnitude is a known property of word embeddings (Schakel and Wilson, 2015). Because ambiguous words appear in diverse, semantically conflicting contexts, their representations are pulled in different directions, reducing their overall magnitude. Unambiguous words, by contrast, carve out a larger, more consistent footprint (and are hence more memorable).

Our account also speaks to two recent studies that used DNN representations to study image memorability, which at first glance appear to be in tension. Ma et al. (2024) found that more memorable images are processed faster in a recurrent CNN – implying memorable stimuli are processed more efficiently. Lin et al. (2024), by contrast, found that images with CNN representations that are harder to reconstruct are more memorable – implying memorable images require more processing. The representational magnitude account reconciles both findings. Images with higher L2 norms activate more features, more strongly; this makes them easier to classify, as the network converges faster when the stimulus aligns well with learned feature detectors. At the same time, stimuli with larger L2 norms are inherently harder to compress, precisely because their representations are more expansive and thus less reducible by a sparse code (indeed, Lin et al. 2024 report a correlation of *ρ >.* 8 between L2 norm and reconstruction error). Both findings are downstream consequences of the same underlying property: a larger representational footprint makes a stimulus easier to classify but harder to reconstruct.

We found a reliable L2 norm effect for visual and lexical memorability, but not for voice memorability. We see two potential explanations for this discrepancy. First, it may be that voice memorability, as compared to memorability of visual stimuli, is itself inherently less consistent across participants, making statistical relations of variables that co-vary with memorability more difficult to observe. Indeed, Revsine et al. (2025) reported that memorability of voices is generally less consistent across individuals than memory for faces, with lower inter-subject agreement (p = 0.28 vs. 0.82) and lower average d’ scores (0.31 for voices vs. 1.11 for faces), which could be largely due to high number of false alarm rates in the voice memorability data. Second, it might be that for voices, there is less alignment between the features driving voice memorability on the one hand, and voice representational magnitude on the other. Based on recent work, voice memorability appears to be primarily driven by relatively lower level acoustic features such as pitch and prosody (Revsine et al., 2025), which separates it from the relatively higher-level features driving visual and lexical memorability, such as category distinctiveness or conceptual/semantic richness (Bainbridge et al., 2017; Xie et al., 2020; Kramer et al., 2023). However, this difference does not explain why we see no consistent relation between voice memorability and representational magnitude, even for early layers of speech models such as Wav2vec2, which should encode such low-level features. It could be that although Wav2vec encodes speaker-specific acoustic information, the dimensions along which it organizes this information do not correspond to the dimensions driving speaker memorability in human perception, such that representational magnitude in Wav2vec does not capture the relevant encoding strength. Characterizing the mechanisms underlying this discrepancy between visual/lexical and voice memorability, and their relationship to representational magnitude, is an exciting avenue for future research.

In this study we focused on recognition memorability, following the original L2 norm effect reported by Jaegle et al. (2019). However, for words, memorability is often not only assessed through a recognition memory test but also through free recall. And while we observed a robust L2 norm effect for word recognition memorability, this effect did not consistently extend to recall memorability. The absence of a consistent effect in recall-based memorability may be due to fundamental differences between recognition and recall processes, which are known to rely on partially distinct neural systems (Cabeza et al. (1997)). Indeed, we found that overall the correlation between lexical recognition and recall memorability was weak (i.e. *r <* 0.1). Moreover, the dataset where this correlation was non-negligible (Cox et al., 2018 with r=0.2; p = 2.18e-1 versus r=0.04 and p = 0.08 in Aka et al., 2023), was also the only dataset in which we observed a relationship between L2 norm and recognition, which further suggests that the representational magnitude effect is primarily related to recognition memory. One possible factor underlying the stronger L2 norm alignment for both recognition and recall memorability in the Cox et al. (2018) dataset might be the task structure. Words were encoded as paired associates, which may have promoted stronger associative encoding but also simultaneously enhanced item-level memory strength for each word, thereby reducing the typical dissociation between recall and recognition. Since recognition may rely more on item-level strength, whereas recall typically depends more on associative and contextual retrieval (Yonelinas et al. (2010)), this task may have shifted recall toward item-based retrieval, making it more similar to recognition. We note, however, that this interpretation remains speculative, as the datasets differ along multiple dimensions, and future work would be needed to test this directly. Our observation that we did not find an L2 norm effect for free recall in two of the three datasets may be explained by the fact that free recall places greater demands on strategic retrieval processes and may be influenced by additional cognitive and contextual factors (Yonelinas et al. (2022)). Additionally, word features such as word frequency seem to affect recognition versus free recall differently: whereas recognition memory tends to be negatively correlated with word frequency (i.e. less frequent words are more easily recognised in an old/new recognition memory test) the opposite tends to be true for free recall: items that are more frequently used in daily language are more easily recalled Aka et al. (2023). Thus, recognition memory versus free recall rely on different cognitive and neural mechanisms which could explain the absence of the representation magnitude effect for recall memorability.

A promising avenue for future research is to directly compare neural representations in the human brain with the representational magnitude of DNNs. While our results show that L2 norm predicts memorability in visual and lexical domains, and aligns with the more general finding that higher response magnitudes during encoding correlates with higher subsequent memorisation rates, it would be interesting to study this systematically, quantifying the joint neural and DNN representational magnitudes for each item. Such studies could subsequently assess whether artificial networks and the brain share similar coding schemes for memorable information, and whether the L2 norm effect is uniform across representational directions or is pronounced for certain ”universal dimensions of representation”, as recently discovered in the visual modality Chen and Bonner (2025).

Together, these results demonstrate that representational magnitude predicts memorability across both images and words, in architecturally distinct networks never designed to model memory. This convergence points to a general principle of distributed representations: the stimuli that make the strongest impression at the representational level – activating the most features, most strongly – are also the stimuli that stick.

## Declaration of competing interest

The authors declare no competing financial interests

## Data and code availability statement

The datasets reported in this paper are all publicly available through the links that are mentioned in the main text. The code used for the analyses is publicly available via: https://github.com/davidamadeusvogelsang/Memorability_L2norm_project.

## CRediT authorship contribution statement

DAV: Conceptualization, data analysis, writing (original+final draft). MH: Conceptualization, writing (original+final draft).

## Declaration of generative AI and AI-assisted technologies in the writing process

During the preparation of this work the author(s) used Chat-GPT4o in order to improve the readiblity of some sentences. After using this tool/service, the author(s) reviewed and edited the content as needed and take full responsibility for the content of the publication.

## Acknowledgements

We thank Adam Osth for his helpful comments and insightful feedback on an earlier version of this manuscript. M.H. is supported by a Veni grant obtained from the NWO (Dutch Research Council)

## Notes

### Competing Interest Statement

The authors have declared no competing interest.

### Summary of Updates

New additions to improve the manuscript based on two expert reviewers in the field

